# Genetic conditionality of strength by the ACE gene in women horse riders

**DOI:** 10.1101/2020.02.12.945444

**Authors:** Lena Majchrowicz, Piotr Matłosz, Rena Majchrowicz

## Abstract

The sport performance relies on hard work, intensity, duration of workout and undoubtedly on genetic background. These components fold on multicausal character of sport achievements. One of the best known genes associated with sport performance is angiotensin converting enzyme gene (ACE). In this study, we showed ACE distribution in women horse riders and the correlation between physiological and genetic background, that should be taken into account when deciding to train sports discipline. Data from 40 healthy female adults were collected. Participants trained horse-riding for at least 3 years and participated in sport competition at least at national level and control group consist of out of training persons. We analysed BMI, %FAT, cardiorespiratory efficiency (VO2max), Complex Reaction Time, body balance, force and Lower Limbs Explosive Force between both calculated groups. Physiological data was calculated by non-parametric Mann-Whitney test with Dunnett’s post hoc test. The DNA amplification from saliva was performed by PCR method to analyze the ACE genotype distribution. In our study, for the first time we showed that the women training dressage have higher force and complex reaction time than the control group. It was correlated with possession of the D allele in genotype that is associated with muscle strength, efficiency and muscle power. There is no research relating to ACE polymorphism among horse riders. This paper showed angiotensin converting enzyme gene distribution polymorphism in female Polish professional horse riders. This article confirms correlation between genotype, sports results and physiological parameters.

## 1. INTRODUCTION

Angiotensin converting enzyme (ACE) is well-known and one of the most frequent under examined gene among sportsmen. An insertion/deletion (I/D) polymorphism in the ACE gene occurrence is responsible for the control of blood pressure in the body [1], as well as for predisposition to short-term or strength sports. It’s been long since known that sport success is the sum of genetic predisposition and regular training. There is a lot of research of frequency of an insertion/deletion of polymorphism among sportspeople training various olympic disciplines. Lack of literature concerning I/D ACE polymorphism with horse-riders [1]. The aim of this study was to show insertion/deletion of polymorphism in the ACE gene correlated with horse-riders’ physical condition and their professing sport discipline and enrich existing literature about genetics of athletic performance. We also wanted to enrich the current canon of knowledge about equestrianism.

Horse-riding is a highly demanding sport rank among olympic disciplines. It split into 7 groups: dressage, show jumping, teaming, western riding, vaulting, long-distance rail and three-day event. Regardless of trained discipline, horse-riding demands high physical condition, force and strength. Riding is most prestigious and well known discipline, but not everyone predispose to this sport [2,3].

All types of sports are marked by determined physical requirements. Sport qualification depend on intensity, length of practice and genetic conditioning of individual facilities. These compounds reflect on the multicasual nature of sport performance [4].

Sport efficiency involves interaction between musculoskeletal, cardiovascular, respiratory and nervous systems. It constitute as one of the most complex properties that predispose to determinated sport. Differences in efficiency can also be the result of anatomy (height, body composition), strength and force. Aerobic efficiency results from the cardiovascular system’s ability to provide and utilize oxygen for muscles. It is not correlated with strength performance, which is related to aerobic threshold [5].

Muscle power is a muscle strength to quantitative force generation. The connection between strength and muscle cramp rate is defined as muscle force. It has a great value in sprint, jumps or weightlifting [6]. Cognitive and enviromental factors (training, nutrition) and injury susceptibility are components of physical fitness. Body response to physical activity partly depends on genetic factors and differs between sports [7]. Physical condition (66%), height (80%) or body build are highly inherited properties connected with force and strength among athletics. Strength and muscle power phenotype research demonstrate 50% aerobic efficiency heritability and 30- 83% strength and force heritability dependent upon type and muscle cramp [4].

Many genes and gene polymorphisms influence on body efficiency and manual effort, such as agene converting angiotensin enzyme ACE [1], ACTN3 α-actinin-3 gene [8], bradykinin β2BDKBR2 receptor gene [9,10], PPAR α receptor gene [11,12], GDF-8 myostatin gene [13,14], APOE apolipoprotein E gene [15]. The ACE gene converting angiotensin I enzyme, which inactivates bradykinin, is part of the renin- angiotensin system responsible for controlling blood pressure in the body. The gene encoding ACE had been localized on 17 chromosomes. Insertion/deletion (I/D) of polymorphism impacts on the predisposition to short-term or strength sports. I/D of polymorphism relies on the presence or absence of a 287 bp Alu repeat element in intron 16 of this gene. The ACE I allele represents an insertion and is associated with lower serum and tissue ACE activity while the D (deleted) allele is associated with higher serum and tissue ACE activity. The ACE I/I genotype is associated with endurance performance and higher exercise efficiency and represents competitive sportspeople and among mountaineers. D/D genotype is associated with strength and power performance. Differences between frequency of I/D ACE polymorphism was observed in population. Abilities to physical effort is also modulated by polymorphism variants in other genes involved in heart and skeletal muscle metabolism [1,4,16].

Most research is concentrated on ACE polymorphism among athletics. Lack of literature concerning I/D polymorphism with horse riders incline us to extend the knowledge about this sport group. The aim of our study was to show the correlation between genetic and physical background at equestrianism.

## 2. MATERIALS AND METHODS

### 2.1. Subject and sample collection

The study was conducted in accordance with the ethical rules of the Helsinki Declaration. Forty people were invited to participate in this study. Eligibility criteria included healthy female adults between the ages of 15 to 35 years old. Participants trained horse-riding (19 persons) for at least 3 years and participated in sport competition at least at national level and control group consist of 21 out of training persons. After explaining the purpose of the study and how to participate, a consent form or permission slip, containing general information about the examined was completed and signed by all participants. The survey was completely anonymous. This study was approved by the University of Rzeszów Bioethics Committee (Poland), under protocol number 2018/01/04.

Briefly, participants expectorated at least 5 mL of unstimulated saliva into a sterile, 20 mL polyethylene tube at least 30 minutes prior to eating, drinking, smoking or kissing to minimize contaminate. Saliva samples were collected from participants and subjected to extract genomic DNA followed by PCR amplification and analysis for ACE I/D polymorphism using specific primers. Collection tubes were maintained on ice and the saliva was aliquoted into sterile 2 mL microcentrifuge tubes containing Saliva DNA Preservative buffer according to manufacturer. Next, each participant’s aliquoted preserved samples were stored at room temperature for three days. The storage periods for the saliva aimed to transport samples into laboratory for further analysis.

### 2.2. DNA extraction

The genomic DNA extraction protocols used are described below. For Saliva DNA Collection, Preservation and Isolation Kit commercial kit (Cat. RU35700, NorgenBiotek Corp., Thorold, Canada) according to the manufacturer’s specific instructions.

Half a milliliter of saliva was collected from each donor into a 2 mL tube (Eppendorf, Hamburg, Germany) containing 0.5 mL Norgen’s Saliva DNA Preservative and mixed by shaking for 10 seconds. In that manner the saliva sample was preserved for storage, shipping and processing. The kit’s procedure was modified by the collection and isolation of saliva DNA from 1 ml of preserved saliva samples, and overnight sample incubation with proteinase K. Isolated DNA was suspended in ddH20. NorgenBiotek commercial kits, the manufacturer’s instructions were followed [17].

### 2.3. Spectrophotometric analysis

The concentrations of DNA from saliva samples were detected using a NanoDrop 2000 (Thermo Fisher Scientific, Waltham, Massachusetts, USA). A spectrophotometer at wavelengths of 230 nm, 260 nm and 280 nm was used to quantify and analyse the condition of each extracted DNA sample from saliva. The concentration and purity of DNA was detected via the relative 260/280 nm and 260/230 nm absorbance ratios. To analyse 1 μL of each sample was required. The 260/280 nm absorbance ratio between 1.6 and 2.0 for samples were considered pure. Ratio lower than 1.6 testifies to higher protein contaminates [17,18].

### 2.4. Electrophoretic analysis

To determine the quality and condition of the extracted DNA, samples from saliva were electrophoresed using a 2% agarose gel in Tris-borate-EDTA (TBE) buffer, Tris base (100 mM), Boric acid (100 mM) and EDTA (2 mM). Standard molecular weight GeneRuler DNA Ladder Mix (SM0333; Thermo Fisher Scientific) in the range of 100 bp to 10,000 bp was loaded onto gel containing Midori Green Advance DNA Stain (MG04; ABO, Gdańsk, Poland). 10 μl of each sample and negative control (10 μl of ddH2O) with 2 μl loading buffer 6× DNA Gel Loading Dye (R0611, Thermo Fisher Scientific) were loaded onto gel. Electrophoresis was driven using 100 V ~ 70 mA for 45 minutes at room temperature. Gels were visualized using UV light and photographed with a digital camera Enduro Gel XL and Transiluminator Gel Vue GVM20 (Syngene, Cambridge, UK).

### 2.5. PCR

Conventional PCR was used to investigate extracted DNA. Polymerase chain reaction detection of the insertion/ deletion polymorphism of human angiotensin converting enzyme gene (ACE). For the I/D of polymorphism, the following primers were used to amplify the region of Alu insertion, with the Alu element’s presence or absence being detected by running the product on an agarose gel. The sequences of primers used were sense oligo: 5’- CTGGAGACCACTCCCATCCTTTCT-3’ (forward) and anti-sense oligo 5’- GATGTGGCCATCACATTCGTCAGAT-3’ (reverse).

Cycling conditions were 95°C for 5 minutes; then for 40 cycles with denaturation at 95°C for 30 seconds, annealing at 62°C for 30 seconds, extension at 72°C for 30 seconds and final extension at 72°C for 10 minutes by Eppendorf Mastercycler personal (Eppendorf). Genotypes of all individuals resulted in an amplified fragment of 190 bp for DD, 490 and 190 bp for ID, and 490 bp for II. The 20 μl reaction mix was composed of 1× GoTaq G2 Green Master Mix (M7823; Promega, Fitchburg, Wisconsin, USA) with dNTPs, MgCl_2_ and Polymerase, 0.1 μM of each primers (upstream, downstream), 100 ng DNA template and nuclease- free water. As a negative control 2 μl nuclease-free water was used. Reaction products were analysed in a 2% agarose gel using the same procedures outlined above (excluding 6× Loading Dye).

### 2.6. Physiological procedures - Body Mass Index (BMI)

To calculate BMI body height and weight was measured in light clothing and barefoot. Body height was measured with SECA (Hamburg, Germany) portable stadiometer to the nearest 0.1 cm according to the protocol recommended by the International Society for the Advancement of Kinanthropometry (ISAK)[19]. Body fat percentage (%FAT) and weight were assessed by bioelectrical impedance using Tanita Body Composition Analyser (TBF-300). BMI was calculated from the equation: BMI =body mass (kg)·stature −1 (m^2^).

### 2.7. Static Balance test

A postural sway was assessed with baropodometric platform (Sensor Medica – FreeMedBase, Rome, Italy). Distribution of the ground reaction forces was used to register Centre Of Pressure (COP) displacements with sampling frequency at the level of 400 Hz. Subjects performed 3 consecutive trials of maximal voluntary forward (FL) and then backward leaning (BL) (Limits Of Stability test - LOS test) proposed by Juras et al. [20]. Subjects stand barefoot on the force platform with feet in natural position, with arms by their sides and palms directed to thighs. Subjects were instructed to look straight ahead on fixation point placed 2 m away on the wall. The procedure started with 10 seconds of quiet standing and then after acoustic signal subjects executed the leaning movement at their own pace until they reached their maximal range. After reaching maximal leaning subjects maintained position to the end of trial (approximately 15 seconds), each trial lasted 30 seconds. Subjects were instructed to execute the leaning only by movement in ankle joint, without raising their heels, or bending in their hip joints or lumbar part of spine. For each trial of forward and backward leaning phase, quantitative measurements of anterior - posterior sway amplitude (ΔX) were assessed to assess limits of stability of each subject.

### 2.8. Lower limbs explosive force, anaerobic power and complex reaction time (RT) assessment

Optojump Next System (Microgate, Bolzano, Italy) was used for lower limbs explosive force, anaerobic power and complex reaction time (RT) assessment. Optojump is an optical measurement system which consist of two parallel bars- one receiver and one transmitter unit. The transmitter bar contains light emitting diodes which communicate continuously with those on the receiving bar. The system detects any interruptions in communication between the bars and calculates their duration measuring thereby flight and contact times during the performance of a series of jumps with an accuracy of 0.001 of a second. Jump height was then estimated as 9.81·flight time^2^·8^−1^ [21]. Bars were placed approximately 1 m apart and parallel to each other. Lower limbs visual and acoustic complex reaction time test involve performance of a jump starting from squad jump position. It was not specified whether the stimulus were to be visual or acoustic. The subject must react to both stimuli. The test result was the average response time of three jumps. Evaluation of lower limbs explosive force involve Counter Movement Jump (CMJ). Subjects perform a single jump starting from an upright position with hands on hips and with counter movement. Subjects were instructed to stand straight up for 1 - 2 seconds and jump as high as it is possible. Single jump started with straight legs and performed a natural flexion before take off. Analysis of anaerobic power involves performance of 15 seconds of jumps, subjects were instructed to perform as many highest jumps as possible during a 15 seconds period. Test result was the average jump power.

### 2.9. Cardiorespiratory efficiency test

Cardiorespiratory fitness was assessed one week apart with aerobic capacity field test - 20 m shuttle run published by Leger et al. [22]. Subjects run as long as possible with continuous movement back and forth between two lines 20 m apart, while keeping the pace with audio signals. The initial speed was 8.5 km·h^−1^ and was increased by 0.5 km·h^−1^ each minute (each stage lasted approximately 1 min). Participants were instructed to run in a straight line, to pivot on completing a shuttle, and to pace themselves in accordance with the audio signals. The subjects were encouraged to keep running as long as possible. The test was finished when the participant failed to reach the end lines concurrent with the audio signals on two consecutive occasions or when the subject stopped because of fatigue. All participants received a comprehensive instruction about the test after which they also practiced the test. They also were instructed to abstain from intense exercises 48 h prior to the test. Tests were carried out under standardized conditions on an outdoor court with hard surface.

The maximal shuttle run speed was used to estimate subjects VO_2max_ using Léger and Gadoury equation for adults: VO_2max_ =−32.678 + 6.592 V, were “V” =last completed stage speed in km·h^−1^ [23].

## 3. RESULTS

### 3.1. Molecular analysis

Performed PCR amplification of DNA from saliva samples showed extensive variability of genotypes in examined people. Insertion allele was identified at size 490 kb and Deletion allele at 190 kb in agarose electrophoresis (Fig.1.).

**Fig.1.**
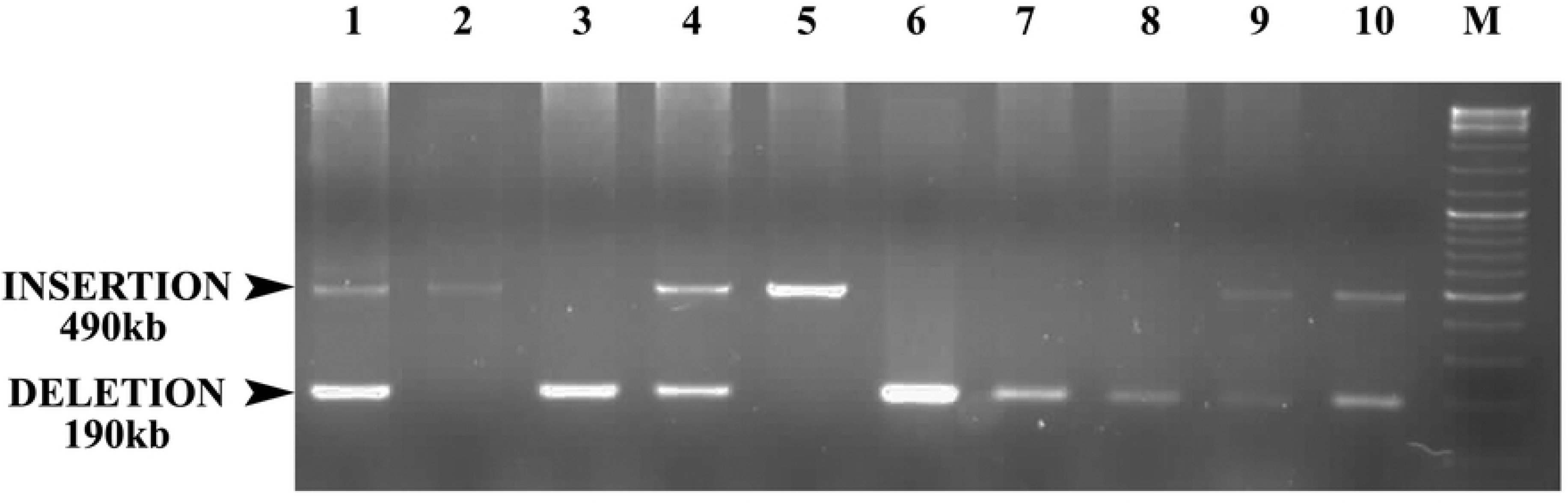
Photography of PCR product showing the ACE genotype distribution among examined people. Lane: 1, 4, 9, 10 - heterozygous I/D; 2, 5 - homozygous I/I; 3, 6, 7, 8 - homozygous D/D; M - DNA ladder

ACE genotype frequency distribution measured by Pearson’s Chi-square (χ^2^) test showed not statistically significant differences between control (CTR) and horse riders (HS) groups. The Chi-square statistic was 1.9973. The p-value was 0.368376. The result was not significant at 0.001> p <0.05. 38% of control group had predisposition to high-performance long-distance sports (I/I genotype), 62% to short-term effort and requiring more muscle strength, efficiency and muscle power (I/D and D/D genotype). Among horse riders sport predispositions were 37% and 63% (respectively) (Fig.2.).

**Fig.2.**
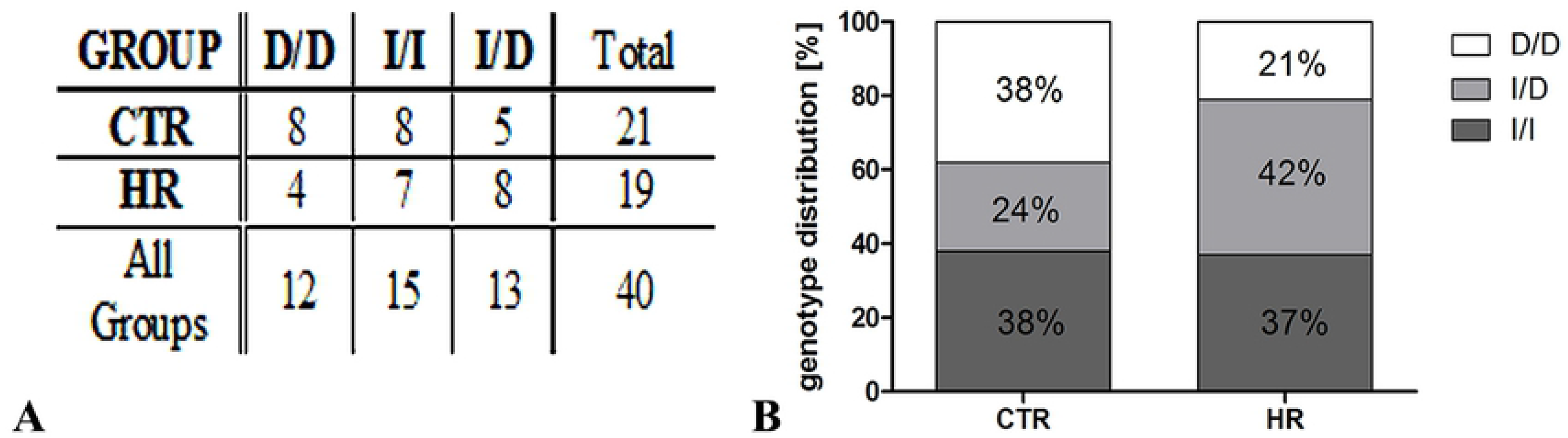
Angiotensin converting enzyme (ACE) genotype frequency distribution measured by Pearson’s Chi-square (χ2) (A) and measured as a percentage (B) in control (CTR) and horse riders (HR) group. Insertion/ Insertion genotype (I/I), Insertion/ Deletion genotype (I/D), Deletion/ Deletion genotype (D/D).

### 3.2. Physiological analysis

Due to the small number of male riders for further analysis we took a group that included only women. We also analysed, BMI, %FAT, cardiorespiratory efficiency (VO_2max_), Complex Reaction Time (CRT), body balance, force and Lower Limbs Explosive Force (LLEF) between both calculated groups. Physiological data was calculated by non-parametric Mann-Whitney test (one-way ANOVA) with Dunnett’s post hoc test. No statistically significant differences were observed between control and horse riders groups in BMI (0.1917), %FAT (0.8927), VO_2max_ (0.4519), CRT (0.1320), balance in posterior (0.1503) and anterior (0.7702) groups and LLEF (0.3971). Only differences at p <0.01 level between force in analysed groups were observed. The p-value was 0.0032.

BMI in all was normal and ranged from 18.2 - 25.7 and 17.1 - 23.8 in control and horse riders group respectively. %FAT in control group was between 13.1 - 30.5% and in examined group was between 10.1 - 31.6%. The higher VO_2max_ (49.72 ml·kg-1·min^−1^) was observed in control group and the lower was 36,54 ml·kg-1·min^−1^. Horse riders were 49.72 ml·kg^−1^·min^−1^ and 33.24 ml·kg^−1^·min^−1^ respectively. Maximum 245.78 mm X axis length (anterior) and 175.66 mm of Y axis length (posterior) and minimum 105.05 mm and 70.97 mm (posterior) were observed in non-training persons. Second group were maximum 223.66 mm and 152.39 mm and minimum 99.68 mm and 69.83 respectively. Complex reaction time oscillated between 0.61 - 0.92 seconds and 0.52 - 1.01 seconds in controls and horse riders group respectively. Lower limbs explosive was between 18.5 - 38.0 cm (CTR) and 19.4 - 34.6 cm (HR). CTR force ranged from 28.97 to 42.2 W·kg^−1^ and HR 25.47 - 61.36 W·kg^−1^.

Moreover, we have classified horse riders into 2 groups. The first one consisted of athletes training jumps, the second were training dressage. We compared every of each groups to control with one-way ANOVA and Dunnett’s a post hoc test. Significant differences were observed only between second study group in force (p <0.01) and CRT (p <0.05) (Fig.3.).

**Fig.3.**
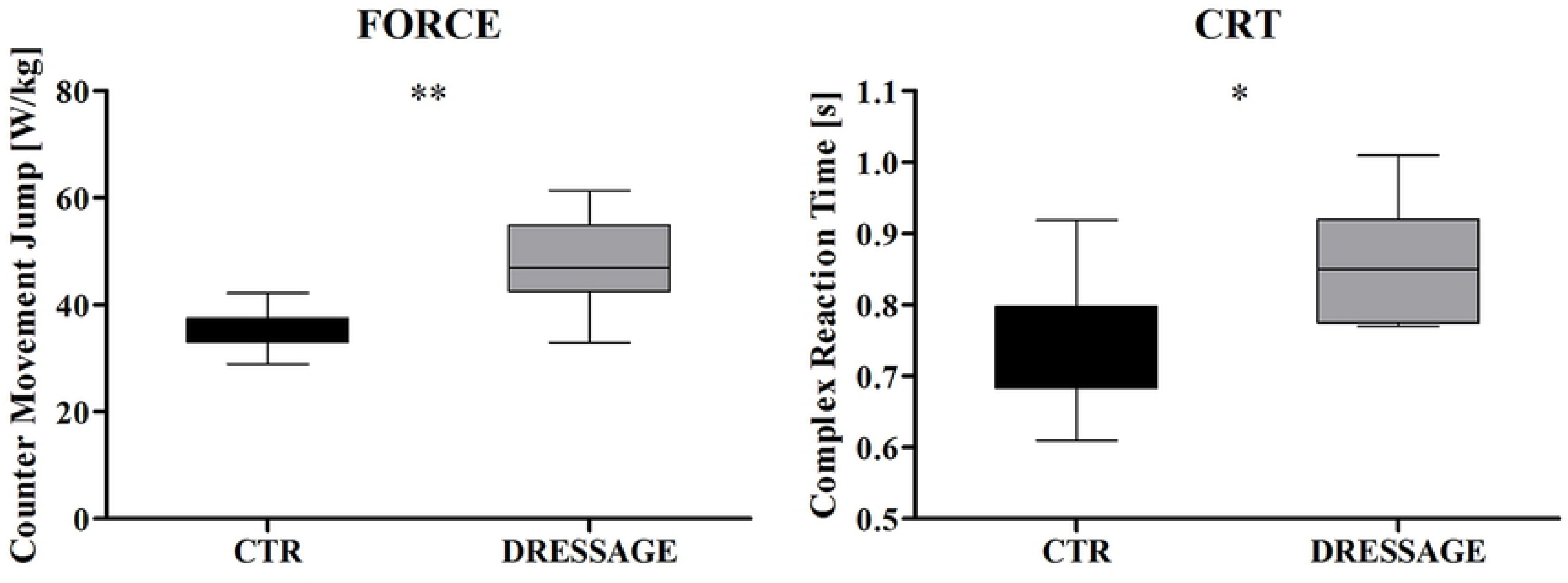
Force and Complex Reaction Time distribution in horse riders training dressage; ***p <0.001, **p <0.01, *p <0.05, no indication - no statistical significance (one-way ANOVA and Dunnett’s a post hoc test). Data presented as mean ± SEM; **P <0.01, by non-parametric Mann-Whitney test.

## 4. DISCUSSION

Polymorphism in angiotensin converting enzyme gene is one of the most frequent examined factors related to sports predispositions. ACE gene is also associated with differences with blood pleasure [24] and fertility [1,25], cardiovascular diseases [1,26–28], neurological diseases like Parkinson’s disease [1,29] or Alzheimer disease [1,30,31]. Since years, many of scientists study the role of Insertion/ Deletion polymorphism and their correlation between genetic and physiological backgrounds in sportspeople. Most research was concentrated on polymorphism in angiotensin converting enzyme (ACE) among athletes [32–36], swimmers [1,37,38], football players [39], volleyball players [24,40] or ball games players [41]. There is no research relating to ACE polymorphism among horse riders.

In this study, we examined angiotensin converting enzyme gene distribution polymorphism in female Polish professional horse riders (jumps and dressage discipline) for the first time. ACE I/I and I/D genotype dominated among examined sportswomen. Moreover, we compared the genomic and physiological results and referred them to control group. D/D and I/I was the most common in control group. Our hypothesis assumed correlation between genotype, sports results and physiological parameters.

We demonstrated that female horse riders especially training dressage have higher force and complex reaction time than sedentary people. These physiological results correlated with possession of D allele in genotype which is associated with muscle strength, efficiency and muscle power.

Genetic background of sport predisposition is relatively easy to verify by molecular biology technics. ACE polymorphisms in combination with physiological tests results, should be taken into account when deciding to train sports discipline in order to achieve success in sport. Nowadays, some medical facilities offer a genetic testing detecting possible polymorphisms within genes: ACTN3, ACE, HIF1A and EPOR what correlate with physiological predispositions to sport.

## 5. CONCLUSION

Our hypothesis assumed correlation between genotype, sports results and physiological parameters. We demonstrated that female HR especially training dressage have higher anaerobic power and CRT than CTR. These physiological results correlated with possession of D allele in genotype which is associated with muscle strength, efficiency and muscle power. In conclusion, ACE polymorphisms combine physiological tests results, should be taken into account when deciding to train sports discipline to achieve success in sport.

## Acknowledgements

The authors would like to thank Katarzyna Zygmunt and PhD Magdalena Gruszk for the help in performing physiological tests, MSc Ewa Banach-Kasper and PhD Maksymilan Bielecki for statistical help.

This work was supported by the University of Rzeszów from the founds of Student Research Society of Diagnostics in Sport and Health Training.

## References

1. Sayed-Tabatabaei FA, Oostra BA, Isaacs A, Van Duijn CM, Witteman JCM. ACE polymorphisms. Circulation Research. 2006. pp. 1123–1133. doi:10.1161/01.RES.0000223145.74217.e7

2. Douglas J-L, Price M, Peters DM. A systematic review of physical fitness, physiological demands and biomechanical performance in equestrian athletes. Comp Exerc Physiol. 2012;8: 53–62. doi:10.3920/CEP12003

3. Gutierrez Rincon JA, Vives Turco J, Muro Martinez I, Casas Vaque I. A comparative study of the metabolic effort expended by horse riders during a jumping competition. Br J Sports Med. 1992;26: 33–35. doi:10.1136/bjsm.26.1.33

4. Guth LM, Roth SM. Genetic influence on athletic performance. Current Opinion in Pediatrics. 2013. pp. 653–658. doi:10.1097/MOP.0b013e3283659087

5. Burnley M, Jones AM. Oxygen uptake kinetics as a determinant of sports performance. Eur J Sport Sci. 2007;7: 63–79. doi:10.1080/17461390701456148

6. Folland JP, Williams AG. Morphological and Neurological Contributions to Increased Strength. Sport Med. 2007;37: 145–168. doi:10.2165/00007256-200737020-00004

7. Puthucheary Z, Skipworth JRA, Rawal J, Loosemore M, Van Someren K, Montgomery HE. Genetic influences in sport and physical performance. Sports Medicine. 2011. pp. 845–859. doi:10.2165/11593200-000000000-00000

8. Yang N, MacArthur DG, Gulbin JP, Hahn AG, Beggs AH, Easteal S, et al. ACTN3 Genotype Is Associated with Human Elite Athletic Performance. Am J Hum Genet. 2003;73: 627–631. doi:10.1086/377590

9. Grenda A, Leońska-Duniec A, Cięszczyk P, Zmijewski P. Bdkrb2 gene −9/+9 polymorphism and swimming performance. Biol Sport. 2014/04/05. 2014;31: 109–113. doi:10.5604/20831862.1096047

10. Saunders CJ, Xenophontos SL, Cariolou MA, Anastassiades LC, Noakes TD, Collins M. The bradykinin β2 receptor (BDKRB2) and endothelial nitric oxide synthase 3 (NOS3) genes and endurance performance during Ironman Triathlons. Hum Mol Genet. 2006;15: 979–987.

11. Lopez-Leon S, Tuvblad C, Forero DA. Sports genetics: the PPARA gene and athletes’ high ability in endurance sports. A systematic review and meta-analysis. Biol Sport. 2015/11/19. 2016;33: 3–6. doi:10.5604/20831862.1180170

12. Tural E, Kara N, Agaoglu SA, Elbistan M, Tasmektepligil MY, Imamoglu O. PPAR-α and PPARGC1A gene variants have strong effects on aerobic performance of Turkish elite endurance athletes. Mol Biol Rep. 2014;41: 5799–5804. doi:10.1007/s11033-014-3453-6

13. Desgeorges MM, Devillard X, Toutain J, Castells J, Divoux D, Arnould DF, et al. Pharmacological inhibition of myostatin improves skeletal muscle mass and function in a mouse model of stroke. Sci Rep. 2017;7: 14000. doi:10.1038/s41598-017-13912-0

14. Elkasrawy MN, Hamrick MW. Myostatin (GDF-8) as a key factor linking muscle mass and bone structure. J Musculoskelet Neuronal Interact. 2010;10: 56–63.

15. Tierney RT, Mansell JL, Higgins M, McDevitt JK, Toone N, Gaughan JP, et al. Apolipoprotein e genotype and concussion in college athletes. Clin J Sport Med. 2010;20: 464–468. doi:10.1097/JSM.0b013e3181fc0a81

16. Dhar S, Ray S, Dutta A, Sengupta B, Chakrabarti S. Polymorphism of ACE gene as the genetic predisposition of coronary artery disease in Eastern India. Indian Heart J. 2012;64: 576–581. doi:10.1016/j.ihj.2012.08.005

17. Garbieri TF, Brozoski DT, Dionísio TJ, Santos CF, Neves LT das. Human DNA extraction from whole saliva that was fresh or stored for 3, 6 or 12 months using five different protocols. J Appl Oral Sci. 2017;25: 147–158. doi:10.1590/1678-77572016-0046

18. Gudiseva H V, Hansen M, Gutierrez L, Collins DW, He J, Verkuil LD, et al. Saliva DNA quality and genotyping efficiency in a predominantly elderly population. BMC Med Genomics. 2016;9: 17. doi:10.1186/s12920-016-0172-y

19. Norton K, Olds T, Olive S, Craig N. Anthropometry and sports performance. Anthropometrica. 1996; 287–364.

20. Juras G, Słomka K, Fredyk A, Sobota G, Bacik B. Evaluation of the Limits of Stability (LOS) Balance Test. J Hum Kinet. 2008;19: 39–52. doi:10.2478/v10078-008-0003-0

21. Bosco C, Luhtanen P, Komi P V. A simple method for measurement of mechanical power in jumping. Eur J Appl Physiol Occup Physiol. 1983;50: 273–282. doi:10.1007/BF00422166

22. Léger LA, Mercier D, Gadoury C, Lambert J. The multistage 20 metre shuttle run test for aerobic fitness. J Sports Sci. 1988;6: 93–101. doi:10.1080/02640418808729800

23. Léger LA, Gadoury C. Validity of the 20 m shuttle run test with 1 min stages to predict VO2max in adults. Can J Sport Sci. 1989;14: 21–26.

24. Durmic TS, Zdravkovic MD, Djelic MN, Gavrilovic TD, Djordjevic Saranovic SA, Plavsic JN, et al. Polymorphisms in ACE and ACTN3 Genes and Blood Pressure Response to Acute Exercise in Elite Male Athletes from Serbia. Tohoku J Exp Med. 2017;243: 311–320. doi:10.1620/tjem.243.311

25. Krege JH, John SWM, Langenbach LL, Hodgin JB, Hagaman JR, Bachman ES, et al. Male◻female differences in fertility and blood pressure in ACE-deficient mice. Nature. 1995;375: 146.

26. de Carvalho SS, Simões e Silva AC, Sabino A de P, Evangelista FCG, Gomes KB, Dusse LMS, et al. Influence of ACE I/D Polymorphism on Circulating Levels of Plasminogen Activator Inhibitor 1, D-Dimer, Ultrasensitive C-Reactive Protein and Transforming Growth Factor β1 in Patients Undergoing Hemodialysis. PLoS One. 2016;11: e0150613.

27. Oudit GY, Crackower MA, Backx PH, Penninger JM. The Role of ACE2 in Cardiovascular Physiology. Trends Cardiovasc Med. 2003;13: 93–101. doi:https://doi.org/10.1016/S1050-1738(02)00233-5

28. Agerholm-Larsen B, Nordestgaard BG, Tybjærg-Hansen A. ACE gene polymorphism in cardiovascular disease: Meta-analyses of small and large studies in whites. Arterioscler Thromb Vasc Biol. 2000;20: 484–492. doi:10.1161/01.ATV.20.2.484

29. Mellick GD, Buchanan DD, McCann SJ, Davis DR, Le Couteur DG, Chan D, et al. The ACE Deletion Polymorphism Is Not Associated with Parkinson’s Disease. Eur Neurol. 1999;41: 103–106. doi:10.1159/000008012

30. Lehmann DJ, Cortina-Borja M, Warden DR, Smith AD, Sleegers K, Prince JA, et al. Large Meta-Analysis Establishes the ACE Insertion-Deletion Polymorphism as a Marker of Alzheimer’s Disease. Am J Epidemiol. 2005;162: 305–317.

31. Kehoe PG, Katzov H, Feuk L, Bennet AM, Johansson B, Wilman B, et al. Haplotypes extending across ACE are associated with Alzheimer’s disease. Hum Mol Genet. 2003;12: 859–867. doi:10.1093/hmg/ddg094

32. Shahmoradi S, Ahmadalipour A, Salehi M. Evaluation of ACE gene I/D polymorphism in Iranian elite athletes. Adv Biomed Res. 2014;3: 207. doi:10.4103/2277-9175.143242

33. Ma F, Yang Y, Li X, Zhou F, Gao C, Li M, et al. The Association of Sport Performance with ACE and ACTN3 Genetic Polymorphisms: A Systematic Review and Meta-Analysis. PLoS One. 2013;8: e54685.

34. Holdys J, Kryściak J, Stanisławski D, Gronek P. ACE I/D Gene Polymorphism in Athletes of Various Sports Disciplines. Hum Mov. 2011;12: 223–231. doi:https://doi.org/10.2478/v10038-011-0022-x

35. Nazarov IB, Woods DR, Montgomery HE, Shneider O V, Kazakov VI, Tomilin N V, et al. The angiotensin converting enzyme I/D polymorphism in Russian athletes. Eur J Hum Genet. 2001;9: 797.

36. Gayagay G, Yu B, Hambly B, Boston T, Hahn A, Celermajer DS, et al. Elite endurance athletes and the ACE I allele - The role of genes in athletic performance. Hum Genet. 1998;103: 48–50. doi:10.1007/s004390050781

37. Tsianos G, Sanders J, Dhamrait S, Humphries S, Grant S, Montgomery H. The ACE gene insertion/deletion polymorphism and elite endurance swimming. Eur J Appl Physiol. 2004;92: 360–362. doi:10.1007/s00421-004-1120-7

38. Woods D, Hickman M, Jamshidi Y, Brull D, Vassiliou V, Jones A, et al. Elite swimmers and the D allele of the ACE I/D polymorphism. Hum Genet. 2001;108: 230–232. doi:10.1007/s004390100466

39. Cięszczyk P, Jastrzębski Z, Zarębska A, Sawczyn M, Drobnik-Kozakiewicz I, Leońska-Duniec A, et al. Association between the ACE I/D polymorphism and physical activity in Polish women. TRENDS Sport Sci. 2016.

40. Ulucan K, Sercan C, Biyikli T. Distribution of Angiotensin-1 Converting Enzyme Insertion/Deletion and α-Actinin-3 Codon 577 Polymorphisms in Turkish Male Soccer Players. Genet Epigenet. 2015;7: 1–4. doi:10.4137/GEG.S31479

41. Jang Y, Kim SM. Influences of the G2350A polymorphism in the ACE Gene on cardiac structure and function of ball game players. J Negat Results Biomed. 2012;11. doi:10.1186/1477-5751-11-6

